# Human first-order tactile neurons can resolve spatial details on the scale of single fingerprint ridges

**DOI:** 10.1101/2020.07.03.185777

**Authors:** Ewa Jarocka, J Andrew Pruszynski, Roland S Johansson

## Abstract

Fast-adapting type 1 (FA-1) and slow-adapting type 1 (SA-1) first-order tactile neurons provide detailed spatiotemporal tactile information when we touch objects with fingertips. The distal axon of these neuron types branches in the skin and innervates many receptor organs associated with fingerprint ridges (Meissner corpuscles and Merkel cell neurite complexes, respectively), resulting in heterogeneous receptive fields that include many highly sensitive zones or ‘subfields’. Using raised dots that tangentially scanned a neuron’s receptive field, here we examined the spatial resolution capacity of FA-1 and SA-1 neurons afforded by their heterogeneous receptive fields and its constancy across scanning speed and direction. We report that the resolution of both neuron types on average corresponds to a spatial period of ∼0.4 mm and provide evidence that a subfield’s spatial selectivity arises because its associated receptor organ measures mechanical events limited to a single fingerprint ridge. Accordingly, the sensitivity topography of a neuron’s receptive fields is quite stable over repeated mappings and over scanning speeds representative of real-world hand use. The sensitivity topography is substantially conserved also for different scanning directions, but the subfields can be relatively displaced by direction-dependent shear deformations of the skin surface.

**Significance Statement:** The branching of the distal axon of first-order tactile neurons with receptor-organs associated with fingerprint ridges (Meissner and Merkel end-organs) results in cutaneous receptive fields composed of several distinct subfields spread across multiple ridges. We show that the spatial selectivity of the subfields typically corresponds to the dimension of the ridges (∼0.4 mm) and that neurons’ subfield layout is well preserved across tangential movement speeds and directions representative of natural use of the fingertips. We submit that the receptor-organ underlying a subfield essentially measures mechanical events at an individual ridge. That neurons receive convergent input from multiple subfields does not preclude the possibility that spatial details can be resolved on the scale of single fingerprint ridges by a population code.

## Introduction

The distal axon of most types of first-order tactile neurons branches in the skin such that a neuron innervates many spatially segregated receptor-organs (Cauna, 1956, 1959; Lindblom and Tapper, 1966; Brown and Iggo, 1967; Jänig, 1971; Looft, 1986; Goldfinger, 1990; Vallbo et al., 1995; Paré et al., 2002; Nolano et al., 2003; Wessberg et al., 2003; Lesniak et al., 2014; Suresh et al., 2016). For the glabrous skin of the human hand, this applies to the fast adapting type 1 (FA-1) and the slowly adapting type 1 (SA-1) neurons which innervate Meissner corpuscles and Merkel cell neurite complexes, respectively (Cauna, 1956, 1959; Vallbo and Johansson, 1984; Nolano et al., 2003), and account for the exquisite tactile spatial acuity of our fingertips (Vallbo and Johansson, 1984). The branching causes the FA-1 and SA-1 neurons to exhibit heterogeneous receptive fields with multiple highly sensitive zones (hereinafter also referred to as subfields) that are seemingly randomly distributed within a circular or elliptical area typically covering five to ten papillary ridges (Johansson, 1978; Phillips et al., 1992; Pruszynski and Johansson, 2014).

We have recently suggested that the subfield arrangement of FA-1 and SA-1 neurons is a functional determinant of the fingertips’ high tactile spatial resolution (Pruszynski and Johansson, 2014; Pruszynski et al., 2018). In short, our hypothesis entails that the spacing between neurons’ interdigitating subfields may constitute the functional limit of the spatial resolution rather than the much greater distance between their receptive field centers as traditionally thought (Johnson and Phillips, 1981; Phillips et al., 1983; Vanboven and Johnson, 1994; Weber et al., 2013). This notion fits well with the fact that the spatial accuracy in the processing of geometric tactile information during object manipulation (Pruszynski et al., 2018) and in certain psychophysical tasks (Loomis and Collins, 1978; Wheat et al., 1995; Dodson et al., 1998; Hollins and Bensmaia, 2007) can exceed that predicted by the Shannon-Nyquist sampling theorem based on the average distance between receptive field centers. However, according to our hypothesis, the intrinsic spatial resolution of the peripheral tactile apparatus would depend not only on the density of the subfields over the skin surface but also on the size of the skin area subtended by each of them, where a smaller size would allow for detection of finer spatial inhomogeneities. Although the spatial resolution of the subfield arrangement of the FA-1 and SA-1 neurons has not yet been quantified in these terms, it seems reasonable to assume that it might approach the dimension of individual fingerprint ridges. First, the receptor organs responsible for the subfields of these neuron types are directly associated with individual papillary ridges (Cauna, 1954; Halata, 1975) and might therefore be able to selectively measure deformations of a single ridge. Second, a ridge can be deflected largely independently of its neighbors (Johansson and LaMotte, 1983; LaMotte and Whitehouse, 1986; Lee et al., 2019).

Here we examined the spatial resolution of the neuron’s subfield arrangement and the robustness of the subfield layout as tactile stimuli slid across the fingertip, as normally occurs during object manipulation and tactile exploration tasks. Unlike with tactile stimuli based on stationary skin indentations, frictional forces can be expected to disrupt neurons’ subfield layout by causing complex direction-dependent shear deformations of the skin within their receptive fields (Johansson and Flanagan, 2009; Delhaye et al., 2016). The receptive field of FA-1 and SA-1 neurons were scanned by a flat surface with small raised dots at speeds representing natural use of our hands in tactile pattern discrimination tasks (15, 30, and 60 mm/s) (Lederman, 1974; Vega-Bermudez et al., 1991; Boundy-Singer et al., 2017; Olczak et al., 2018). We quantified the spatial resolution of the receptive fields’ subfield arrangement and examined how robustly a neuron’s subfield layout spatially structures its response in the space domain across repeated scans and across scanning speeds. We also compared neurons’ subfield resolution and the spatial structuring of their responses when scanning the receptive field in two opposite tangential directions, equivalent to moving our fingertips back and forth while exploring tactile surfaces.

## Materials and Methods

### Participants and general procedure

Twelve healthy humans, 20–30 years of age (6 females), participated after providing written informed consent in accordance with the Declaration of Helsinki. The Umeå University ethics committee approved the study.

Each subject reclined comfortably in a dentist’s chair with the right upper arm abducted ∼30°, the elbow extended to ∼120°, and the hand supinated. A vacuum cast supported by a metal frame immobilized the forearm, and Velcro strips around the wrist provided additional fixation. To stabilize fingertips, we glued the nails to plastic holders firmly attached to the frame that also supported a robot that controlled the tactile stimulation (Birznieks et al., 2001).

Action potentials from single first-order tactile neurons terminating in the glabrous skin of the distal segment of the index, long or ring finger were recorded with tungsten electrodes (Vallbo and Hagbarth, 1968) percutaneously inserted into the right median nerve at the mid-level of the upper arm. Isolated neurons were classified as fast-adapting type 1 (FA-1), slow-adapting type 1 (SA-1), fast-adapting type 2 (FA-2), and slow-adapting type 2 (SA-2), according to previously described criteria (Vallbo and Johansson, 1984). We focused on FA-1 neurons (N = 23) and SA-1 neurons (N = 11) whose well-defined cutaneous receptive fields are made up of a number of subfields (Johansson, 1978; Phillips et al., 1992; Pruszynski and Johansson, 2014).

### Tactile stimuli

We analyzed the neurons’ responses elicited by a stimulus pattern that contained raised dots on a flat background moving tangentially across the receptive field and (**Fig. 1A**). The dots were 0.45 mm high truncated cones with a flat 0.4 mm diameter top and a base diameter of 0.7 mm (**Fig. 1B**, see inset). The stimulus pattern was produced via a standard photo-etching technique using a photosensitive nylon polymer (Toyobo EF 70 GB, Toyobo Co., Ltd., Osaka, Japan) and wrapped around a transparent rotating drum (**Fig. 1B**). A custom-built robotic device controlled the rotation speed of the drum and kept the normal contact force between the stimulus pattern and the receptor-bearing fingertip constant at ∼0.4 N (for details see (Pruszynski and Johansson, 2014). This force was chosen because it conveniently falls within the range that humans use to manually explore surfaces (Lederman, 1974; Gamzu and Ahissar, 2001; Smith et al., 2002; Olczak et al., 2018). A video camera mounted in the transparent drum was used to position the stimulus pattern with reference to the location of the neuron’s receptive field on the fingertip, outlined with an ink pen as previously described (Johansson and Vallbo, 1980).

**Figure 1.**
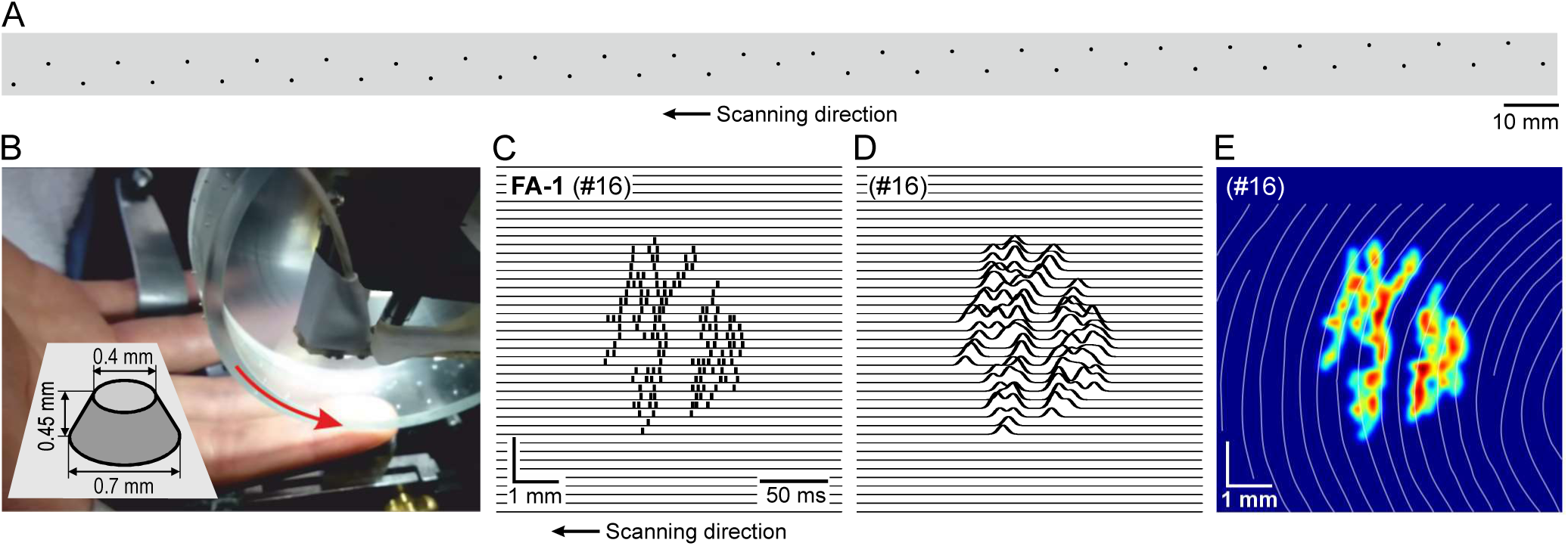
Experimental setup, stimuli and basic approach. Stimulating surface with raised dots for mapping receptive field sensitivity topography of first-order tactile neurons. **B**. The surface was wrapped around a transparent drum and a custom-built robotic device controlled the drum’s rotation and position. Inset shows schematically one of the raised dots. **C**. Two-dimensional spatial event plot for an exemplar FA-1 neuron (#16) obtained during one drum rotation in proximal-distal direction at 30 mm/s tangential speed. **D**. Sensitivity map obtained after convolving spike events in **C** with a one-dimensional kernel (SD = 0.1 mm). **E**. Color coded sensitivity map obtained after convolving the same spike events with a two-dimensional kernel (SD = 0.1 mm). The white lines mark the grooves between the papillary ridges.

The stimulus surface included stimuli used to generate a sensitivity map of the neuron’s receptive field (**Fig. 1A**). The layout of the field mapping dots was designed to generate a field sensitivity map for each drum revolution based on one dot stimulating the neuron at the time. Forty-one dots were equally distributed along the extent of the stimulation surface in the movement direction (length = 312 mm), defined as the x-direction. In the perpendicular direction, defined as the y-direction, the dots were equally spaced on the 8 mm wide zone. Thus, the dots moved over the skin in separate tracks spaced 0.2 mm apart. Overall the dots were spaced at least 7 mm apart to minimize interactions between neighboring dots on a neuron’s response (Phillips et al., 1992). The array contained four additional dots that were evenly spaced in the x-direction and located in the center of the mapping zone in the y-direction. These dots, which nominally moved across the center of the neuron’s receptive field, could be used to control the spatial alignment of action potentials in the movement direction.

### Stimulation protocol

For each neuron, the field mapping zone was moved over its receptive field in the proximal-distal direction of the finger at a speed of 30 mm/s for four drum revolutions. Thereafter, the drum was laterally repositioned to expose the receptive field to another stimulation pattern containing raised elements for 15 drum revolutions in the same direction and movement speed (data not shown). For neurons with stable enough recording of unitary action potentials, the corresponding protocol was then run at a speed of 60 mm/s and at 15 mm/s (10 FA-1s and 6 SA-1s). For neurons with still discriminable unitary action potentials, we then ran the above scheme with drum rotations at 30 mm/s in the opposite direction, i.e., the stimulus pattern now moved in the distal-proximal direction (8 FA-1s and 3 SA-1s). For analysis of the effect of scanning direction on receptive field sensitivity topography we also included data for proximal-distal and distal-proximal stimulation at 30 mm/s gathered in a previous series of experiments but not analyzed direction-wise (8 FA-1s and 3 SA-1s) (Pruszynski and Johansson, 2014).

### Data processing and analysis

The nerve signal, the instantaneous position of the stimulus surface recorded via a drum shaft decoder (AC36, Hengstler GmbH) providing a resolution of 3 μm and the contact force were digitally sampled at 19.2 kHz, 2.4 and 0.6 kHz respectively (SC/ZOOM, Department of Integrative Medical Biology, Umeå University). Unitary action potentials were detected on-line based on spike morphology and verified for each action potential off-line (Edin et al., 1988).

For each drum revolution, we constructed a two-dimensional spatial event plot (SEP (Johnson and Lamb, 1981)) of the neuron’s receptive field based on the position of the stimulating dot at each evoked action potential (**Fig. 1C)**. Since the dots were distributed along the direction of motion of the stimulus pattern (x-direction), the instantaneous x-coordinate of the stimulating dot was offset based on its known x-coordinate. The y-position was defined by the y-coordinate of the track (one out of 41) in which the stimulating dot moved. In the analysis, data from the first of the four drum revolutions were omitted and the second revolution was used as the first scan of the receptive field. We took this action to minimize distorting effect of creep deformation of the fingertip on construction of SEPs since visual inspection of the skin revealed that the major creep deformation occurred during the first revolution. To render a smooth receptive field map, we convolved with a Gaussian function SEPs obtained from each drum revolution within an 8 by 8 mm window centered on the centroid of the spike activity. **Figure 1D** illustrates a receptive field sensitivity map obtained by convolving a neuron’s spike traces with a kernel width of 0.1 mm and **Fig. 1E** shows a color-coded map generated with a corresponding two-dimensional Gaussian where brighter colors indicate higher spatial density of action potentials. For each SEP, mean firing rate was calculated as the number of spikes evoked within the 8 by 8 mm window divided by the duration of the stimulus dot within this window. Peak firing rate was defined as the reciprocal of the shortest, stimulus-evoked interspike interval observed in a SEP.

Our estimates of a neuron’s spatial resolution and how its receptive field sensitivity topography can be affected by scanning speed and direction relied on assessment of similarity between maps based on pairwise correlations of SEPs after convolution with Gaussians of different widths defined in the spatial domain. By convolving each SEP with 21 different kernels with logarithmically spaced standard deviations in the range of 0.02 to 0.33 mm we gradually simulated increased noise on the positions of the action potentials, which increasingly blurred the representation of the field sensitivity topography. Our approach is analogous to methods previously used to assess the similarity of pairs of individual spike trains obtained under different experimental conditions but when represented in the temporal domain (Schreiber et al., 2003; Fellous et al., 2004; Vazquez et al., 2013; Pruszynski and Johansson, 2014).

To account for skin warping in the analysis of effects of scanning direction on the receptive field sensitivity topography, we used iterative cross-correlation to define parameters for transforming the map obtained in the distal-proximal scanning direction to best resemble that obtained in the proximal-distal direction. This implied finding the parameter values for obtaining the maximum correlation coefficient after ± 4 mm stretching/compression of the map (8 × 8 mm) in increments of 0.2 mm in the scanning direction and in its perpendicular direction, and rotation of the map ± 20° in 2° increments, taking into account every possible combination in the matrix.

To relate a neuron’s receptive field sensitivity topography to the layout of the fingerprint ridges, we overlaid the sensitivity map on manual tracings of the papillary grooves of the appropriate skin surface based on a still image taken from the recorded video; the spatial acuity of our video surveillance system and its temporal resolution (1 frame per 40 ms) were not sufficient for analysis of time-varying correlations between spike events and deformation changes of individual ridges caused by dot the stimulus. We calculated an average of the ridge width (RW) in the receptive field by measuring the length of a line oriented such that it transversely crossed 5 ridges centrally in the field. We also recorded the orientation of this line with reference to the scanning direction (α). Although the basic types of fingerprints are arches, radial loops, ulnar loops and whorls, when a small area corresponding to the size of the current receptive fields is considered, the ridges were usually quite parallel (see Results). We estimated the distance by which the leading edge of the dot stimulus travelled across a ridge by considering the ridge orientation relative to the scanning direction. The shortest distance occurs when a ridge is oriented perpendicular to the scanning direction, while the distance gradually increases when the ridge orientation becomes increasingly oblique relative to the scanning direction. The increase of the distance as a function α was calculated as RW/cosine(α)-RW.

### Experimental Design and Statistical Analysis

Effects of the experimental factors on neural response variables were assessed using two-tailed t-test for independent samples by groups and two-way mixed-design ANOVAs with neuron type (FA-1 and SA-1) as a between-group effect. We used the Tukey HSD test for post-hoc comparisons. Correlation coefficients were Fisher transformed into Z scores when performing parametric statistics and in estimating average values that were then converted back to correlation coefficients. Correlation values are reported as coefficients of determination (R^2^). All statistical tests were deemed significant if P < 0.05. Unless otherwise stated, reported point estimates based on sample data refer to mean ±1 standard deviation (SD).

## Results

We present the results in four sections. First, we estimate the spatial resolution of FA-1 and SA-1 neurons with respect to the receptive field’s internal sensitivity topography and provide evidence suggesting that the resolution of a subfield matches the dimension of an individual fingerprint ridge. Second, we analyze the similarity of a neuron’s receptive field maps obtained across repeated mappings and address heterogeneity amongst neurons regarding the subfield layout. Third, we test how well a neuron’s field sensitivity topography is maintained at different scanning speeds (15, 30 and 60 mm/s). Fourth, we investigate the consistency of the receptive sensitivity topography across different stimulation directions by comparing results from scans in the proximal-distal and distal-proximal direction.

### Spatial resolution of subfields

To estimate a neuron’s spatial resolution with respect to the receptive field’s internal sensitivity topography, we first generated a set of receptive field maps by convolving the spatial event plot (SEP) obtained at each scan with a two-dimensional Gaussian function at 21 different kernel widths with standard deviations increasing from 0.02 to 0.33 mm (**Figs. 2A, C**). Thus, we simulated gradually increased noise on the positions of the action potentials, which increasingly blurred the representation of the sensitivity topography of the receptive field (**Figs. 2B, D**). We then calculated the pairwise two-dimensional cross-correlation between the three maps which resulted in 3 correlations per kernel width and stimulation condition (scanning speed and direction) (**Figs. 2B, D**).

**Figure 2.**
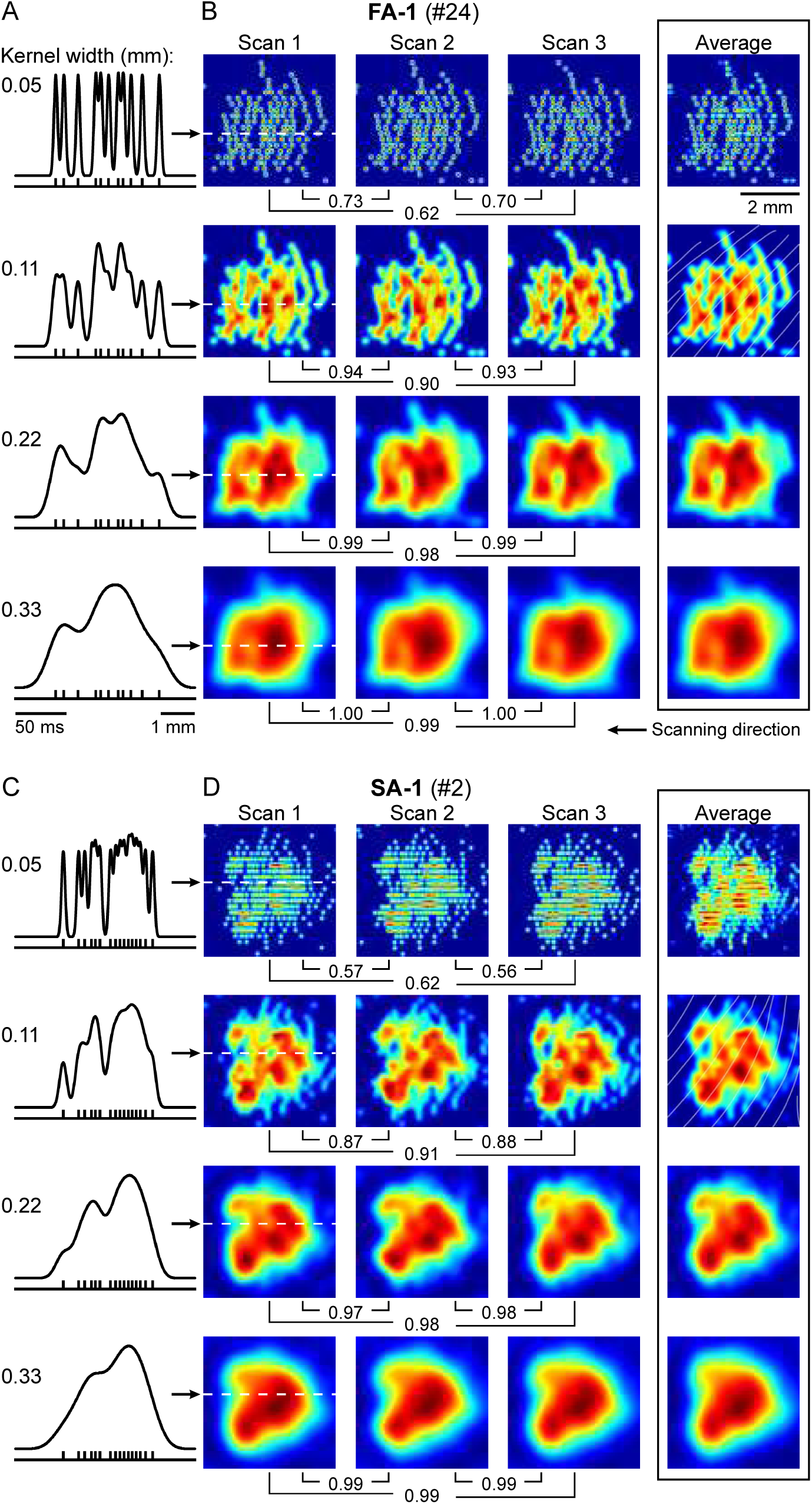
Effect of kernel width on the receptive field map. **A**. The different panels show, for an exemplar FA-1 neuron (#24), one spike train elicited by one of the stimulus dots when passing along its track over the receptive field (dashed white line in B) and this train convolved with four of the 21 different kernels used (SD = 0.05, 0.11, 0.22 and 0.33 mm; top trace). **B**. Sensitivity maps of the same neuron obtained by convolving the spatial event plots generated during each of the three scans (Scan 1 – 3) with the kernel widths shown in A. The rightmost sensitivity maps represent, for each kernel width, the average of the three maps. The numbers indicate R^2^ values of correlated pairs of maps. The white lines mark the grooves between the papillary ridges. **C** – **D**. Data from an exemplary SA-1 neuron (#2) shown in the same format as in A and B. **A** – **D**. Neurons scanned at 30 mm/s in the proximal-distal direction.

As illustrated in **Fig. 3A**, for all 34 neurons stimulated at 30 mm/s in the proximal-distal direction the correlation between the three maps increased as a function of kernel width (solid lines). The low correlations obtained with the narrowest kernels were caused by the spike jitter between repetitions of the same stimulus tending to be greater than the kernel width. With gradually wider kernels, the correlation quite steeply increased up to around 0.1 mm width and then remained high as the maps became more Gaussian-shaped and moved towards having a single point of maximum sensitivity (**Figs. 2B, D**). The kernel width at this breaking point (∼ 0.1 mm) offered an initial estimate of the spatial resolution of the neurons’ sub-field arrangement since additional spatial filtering that attenuated the sensitivity topography of the receptive field did not substantially increase the correlation. To further assess the reliability of this estimate, we compared the mean value of the correlations between the three empirical maps as a function of kernel width with the corresponding mean of pairwise correlations between each of the three empirical maps and the same map rotated 180° (3 correlations for each kernel width). Through the rotation, we confused the internal sensitivity topography of a neuron’s receptive field while maintaining its generally oval shape, its orientation and size. As expected, the correlation involving map rotation increased more slowly with increasing kernel width, especially for widths below ∼0.1 mm (**Fig. 3A**, dashed lines). We calculated the difference between the mean values for the correlations between the empirical maps and the correlations that included 180° map rotation as a function of the kernel width and used the kernel width where the difference was maximal as a point estimate of a neuron’s spatial resolution (**Figs. 3B, C**). For neurons scanned at 30 mm/s in the proximal-distal direction, the estimated resolution was on average 0.081 ± 0.025 mm (mean ± 1SD, N = 34) and did not reliably differ between neuron type (t_32_ = 0.73; P = 0.47; t-test for independent samples by groups).

**Figure 3.**
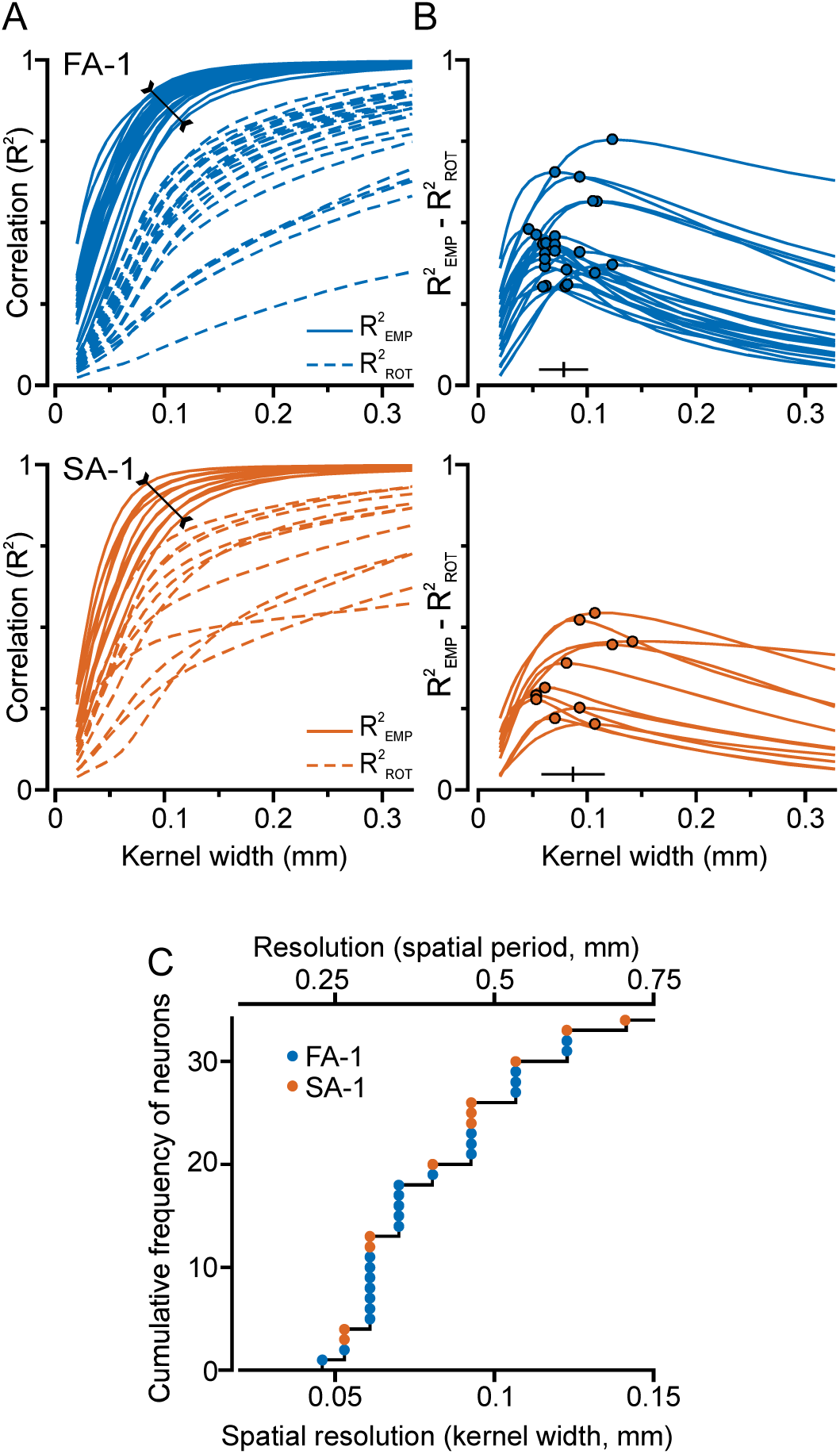
Spatial resolution when scanned at 30 mm/s in the proximal-distal direction. **A**. Superimposed curves show, for individual neurons (23 FA-1s, 11 SA-1s), mean values of pairwise correlations between the empirical maps obtained during the three scans (solid lines, R^2^ _EMP_) and of correlations between each of the three empirical maps and the same map rotated by 180° (dashed lines, R^2^ _ROT_) as a function of kernel width. The slanted line with arrowheads at the ends, centered on 0.1 mm kernel width, roughly marks the breaking point where further spatial filtering did not substantially increase the correlations between the empirical maps. **B**. Difference between correlations amongst the empirical maps and those involving 180° map rotation for individual neurons as a function of kernel width. The filled circles indicate the point of maximum difference for each neuron and the horizontal bar indicates mean ±1 SD across neurons of the kernel width at this point. **C**. Distribution across neurons of the estimated spatial resolution represented as the kernel width yielding the maximum difference (bottom abscissa) and as a sinusoidal spatial period (top abscissa). Clustering of data points at different abscissa values results from the kernel widths used for convolving with the spike trains (see Methods).

Given that the receptor organs of FA-1 and SA-1 neurons are associated with individual fingerprint ridges, we sought to relate a neuron’s resolution to the dimensions of the ridges within its receptive field. For this, we expressed a neuron’s subfield resolution as a sinusoidal spatial period by utilizing the fact that a basic cosine cycle specified between –π and π is very similar to a Gaussian function within ± 2.5 SDs (R^2^ = 0.996). Hence, in sinusoidal terms, the resolution was ∼ 0.41 ± 0.12 mm averaged across the neurons (i.e., 5 times that expressed as kernel width; upper abscissa in **Fig. 3C**). Measurements within the neurons’ receptive fields indicated that the width of the fingerprint ridges was similar: 0.47 ± 0.10 mm (N = 33; video image was missing for one SA-1 neuron). These results suggested that the spatial selectivity of the subfields basically matched the width of an individual ridge. Likewise, inspection of the receptive field maps gave the impression that the dimension of individual subfields often corresponded to the width of a ridge (**Fig. 1E**, also see **Fig. 5B**).

**Figure 4.**
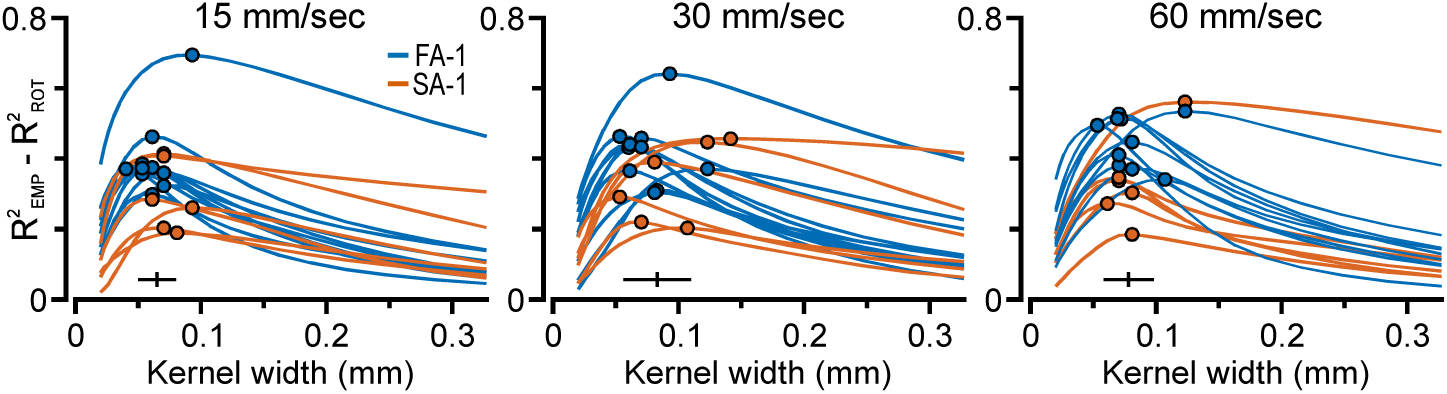
Spatial resolution at different scanning speeds. Difference between correlations amongst the empirical maps and those involving 180° map rotation for individual neurons mapped at all three scanning speeds (15, 30 and 60 mm/s) as a function of kernel width. Same format as Fig. 3B.

**Figure 5.**
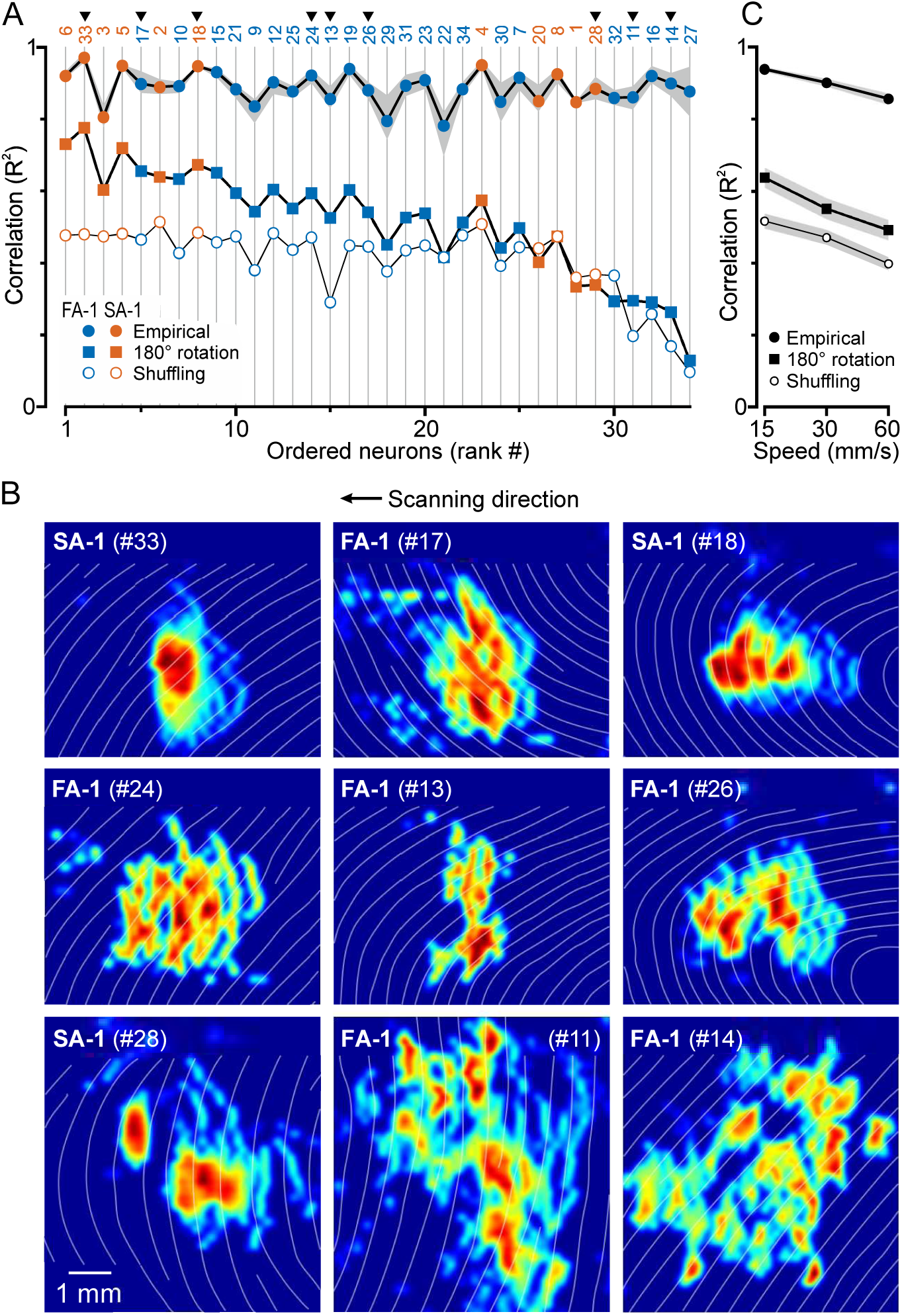
Consistency and heterogeneity of neurons’ subfield layout. **A**. Filled circles joined by the top curve show, for each neuron (23 FA-1s, 11 SA-1s), the mean value of the three correlations between the maps of the three scans at 30 mm/s in the proximal-distal direction. The gray shading indicates the range of these correlations. Correspondingly, filled squares show the mean value of the correlations involving 180° map rotation, and hollow circles show the mean correlation between each of the three empirical maps and each of the corresponding maps of all other neurons (“shuffling”). Numbers at the top indicate the identification number for each neuron used throughout the paper and arrowheads indicate neurons featured in B. Neurons have been ranked along the abscissa as a function of increasing difference between the correlations amongst the empirical maps and those involving 180° map rotation. **B**. Examples of sensitivity maps of neurons with small, intermediate and large difference (top, middle and bottom panels, respectively) obtained by scans at 30 mm/s in the proximal-distal direction; average map across the three scans is shown. The white lines indicate the grooves between the papillary ridges. **C**. Mean R^2^ values from A as function of scanning speed (15, 30 and 60 mm/s). The gray shading indicates standard error of the mean (N = 16).

We asked if the spatial selectivity of a neuron’s subfields is directly linked to the width of the ridges in its receptive field by utilizing the variability between neurons in these parameters in multiple linear regression with the neurons’ estimated spatial resolution as the dependent variable and with ridge width as one independent variable. Since the orientation of the ridges relative to the scanning direction could vary between neurons (see **Figs. 5B** and **6A**), a second independent variable dealt with the possibility that the subfields had a farther extent and thus a poorer resolution when the stimulus moved along the ridges compared with mainly across the ridges. Specifically, this variable indicated the increase in distance that the stimulus interacted with the ridges depending on the obliqueness of their orientation relative to the scanning direction (see Methods). A reliable regression equation was found (R2 = 0.27, F_(2, 30)_ = 5.67, P = 0.008). Both, ridge width and distance increase were significant predictors of spatial resolution (β = 0.49, P = 0.005 and β = 0.38, P = 0.027, respectively). The predicted resolution expressed as spatial period was equal to 0.08 + 0.63 x (ridge width) + 0.08 x (increased distance), all measures in mm. Thus, the spatial period representing a neuron’s resolution increased by 0.063 mm for each 0.1 mm increase in ridge width. However, it only increased by 0.008 mm for each 0.1 mm increase in the stimulation distance along the ridges, suggesting that the direction of stimulation with reference to ridge orientation had a relatively limited influence on the resolution of the subfield arrangement. Overall, these results are consistent with the idea that a subfield essentially records tactile events localized to a limited segment of an individual ridge.

The effect of scanning speed (15, 30 and 60 mm/s) on the spatial resolution was investigated for 10 FA-1 and 6 SA-1 neurons stimulated in the proximal-distal direction. For the 15 and the 60 mm/s scanning speeds, the effect of kernel width on the correlations between the empirical maps and those involving 180° map rotation maps was similar to that for 30 mm/s (**Fig. 4**). We found an effect of speed on the spatial resolution (F_2,28_ = 5.00, P = 0.014), the kernel width tended to be smaller at 15 mm/s (0.065 ± 0.015 mm) than at 30 mm/s (0.083 ± 0.027 mm; P = 0.01, Tukey HSD post-hoc test) and 60 mm/s (0.078 ± 0.020 mm; P = 0.08) and did not statistically differ between 30 and 60 mm/s (P = 0.63). There was no effect of neuron type (F_1,14_ = 0.92 P = 0.35) and no interaction effect between speed and neuron type (F_2,28_ = 1.35, P = 0.27).

In sum, the spatial resolution of the subfield arrangement of the FA-1 and the SA-1 neurons corresponded to kernel widths around 0.1 mm and slightly below, it was barely affected by scanning speed and expressed as spatial period it matched the width of single fingerprint ridges. The remainder of the results section is based on analyses where we consistently used receptive field maps obtained with a kernel width of 0.1 mm. Note that none of our conclusions were qualitatively altered with corresponding analyzes based on kernel widths identified for each individual neuron.

### Consistency and heterogeneity of neurons’ subfield arrangement

In this section, we analyze the similarity of a neuron’s receptive field maps obtained from the repeated mappings in the proximal-distal direction and address heterogeneity of neurons’ receptive field sensitivity profile.

A neuron’s receptive field maps obtained at the three consecutive scans at a given speed and direction were very similar (see **Fig. 2B, D)**. For scans at 30 mm/s, the mean correlation for the three pairwise cross-correlations obtained for the individual neurons (**Fig. 5A**, filled circles) averaged 0.90 (mean R^2^; median = 0.89) across the 34 neurons and did not differ reliably between neuron type (t_32_ = 1.95; P = 0.06; t-test for independent samples by groups). The variability in R^2^ values across the pairwise correlations was small (**Fig. 5A**, gray area around top curve).

To provide a reference for the correlation observed across repeated mappings with regard to the significance of the subfield layout, first we used the pairwise correlations between each of the empirical maps and the same map rotated by 180° (**Fig. 5A**, filled squares). As indicated above, these correlations involved disruption of the subfield layout while being modestly affected by the oval overall shape of the receptive field and its orientation. Second, we cross-correlated each of the three empirical maps with each of the maps obtained at the corresponding scan of all other neurons (3 × 33 = 99 correlations per neuron; **Fig. 5A**, open circles). This “shuffling” would yield correlations that might be worse because they would also be sensitive to the principal orientation as well as to the overall size of the receptive field.

In **Fig. 5A**, the neurons are ranked along the abscissa based on the difference between the correlation of the empirical maps and the correlation involving 180° map rotation. Averaged across all neurons, the latter correlation was markedly lower than the correlation between the empirical maps (mean R^2^ = 0.52 vs. 0.90). However, the difference varied substantially between neurons (vertical distance between the filled circle and squares in **Fig. 5A**). Neurons with the smallest differences (subfield arrangement least sensitive to receptive field rotation), usually had quite complex receptive fields but with a noticeable 180° rotational symmetry, or occasionally a field with essentially only one highly sensitive zone (**Fig. 5B**, top row). Neurons with intermediate differences usually showed complex multifocal receptive fields (**Fig. 5B**, middle row) and those with the largest difference typically had very patchy receptive fields with widely spread subfields (**Fig. 5B**, bottom row). A two-way mixed design ANOVA applied to the difference in the correlations involving the empirical maps and those involving map manipulations (180° rotation, shuffling) indicated a main effect of the map manipulation (F_1,32_ = 38.39, P < 0.0001) but not of neuron type (F_1,32_ = 0.23, P = 0.63) and no significant interaction effect (F_1,32_= 0.55, P = 0.46). The field shuffling yielded a weaker correlation than the 180° rotation. The differential effect of the 180° rotation and the shuffling could markedly vary between neurons (**Fig. 5A**) where neurons with widely scattered subfields were similarly affected.

For the 16 neurons scanned at all three speeds (15, 30 and 60 mm/s) in the proximal-distal direction, scanning speed affected the correlations between the empirical maps (F_2,28_ = 58.76, P < 0.0001). The average R^2^ of the empirical maps was 0.94, 0.90 and 0.86 at 15, 30 and 60 mm/s, respectively (**Fig. 5C**). There was no main effect of neuron type (F_1,14_ = 0.09, P = 0.77) or interaction effect between speed and neuron type (F_2,28_ = 1.29, P = 0.29). As with 30 mm/s, the variability in the pairwise correlations at 15 and 60 mm/s was small. For 15 and 60 mm/s, the effect of the map manipulations was like that described above for 30 mm/s (**Fig. 5C**). That is, for the neurons scanned at all three speeds, a three-way mixed design ANOVA failed to indicate an effect of speed and neuron type on the difference in mean correlations between the empirical maps and the correlations involving the field manipulations (F_2.28_ = 2.06, P = 0.16 and F_1.14_ = 1.94, P = 0.18, respectively), while map manipulation had a main effect (F_1.14_ = 39.00, P <0.0001) with a greater difference with shuffling than with 180° map rotation. There were no significant interaction effects between these factors.

In sum, these results show that the sensitivity topography of the receptive fields is well conserved across consecutive scans regardless of speed but can be quite heterogeneous across neurons.

### Conservation of receptive field sensitivity topography across scanning speeds

Based on data from neurons scanned at all three speeds in the proximal-distal direction, we asked to what extent a neuron’s subfield layout is maintained across scanning speeds. In this analysis we used an average of the three maps obtained with each scanning speed constructed with the 0.1 mm kernel width.

Visual inspection of the maps indicated that a neuron’s subfield stood out with a similar layout at all speeds (**Fig. 6A**). However, decreases in speed resulted in an increased maximum spike density in the subfields, which is consistent with previous results regarding the effect of speed on the number of action potentials of a neuron’s spatial event plot (Phillips et al., 1992). A two-way ANOVA verified that speed influenced the number of action potentials (F_2,28_ = 61.5, P < 0.0001) but not neuron type (F_1,14_ ≤ 2.55, P ≥ 0.13). Considering firing rates, the mean as well as the peak firing rate increased with increasing speed (F_2,28_ = 88.9, P < 0.0001; F_2,28_ = 17.4, P < 0.0001, respectively), with no significant effect of neuron type (F_1,14_ ≤ 2.55, P ≥ 0.13 in both instances). Averaged across the three scans and all 16 neurons, the mean rate was 11 ± 5, 21 ± 7, 26 ± 9 Hz (mean ± 1SD) at 15, 30 and 60 mm/s, respectively. With increasing speed, the spikes were generated for shorter periods, yet the mean firing rate did not increase proportionate to speed because of the decreasing ratio of number of spikes per scan to speed. For the peak rate, the speed effect was modest. Averaged across all 16 neurons, the peak rate was 210 ± 53, 245 ± 66, 244 ± 53 Hz at 15, 30 and 60 mm/s, respectively.

**Figure 6.**
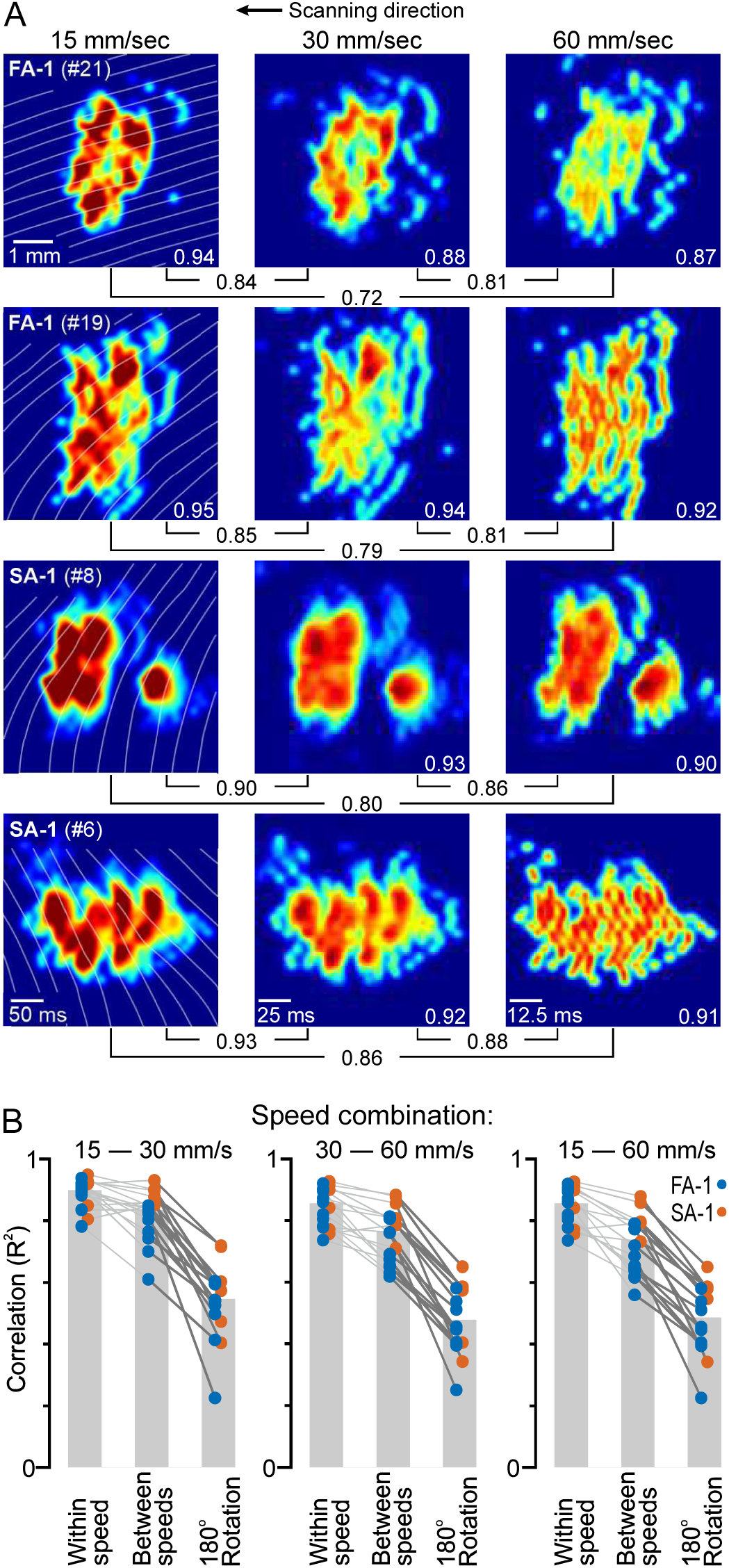
Effect of scanning speed on sensitivity topography. **A.** Sensitivity maps obtained at 15, 30 and 60 mm/s scanning speed for two exemplar neurons of each type (FA-1 and SA-1); average map across the three scans is shown. The white lines mark the grooves between the papillary ridges. The numbers underneath the maps indicate R^2^-values of correlated pairs of maps (between-speed correlations) and the number on the maps indicate, for each map, the mean R^2^ of the correlations between the empirical maps obtained during the three scans (within-speed correlations). **B**. For each speed combination and neuron, symbols show: (i) average correlation between the three empirical maps for the speed that showed the lowest correlation (“Within speeds”); (ii) pairwise correlation values between average maps obtained with the respective speed (“Between speeds”); (iii) average correlation between each of the empirical maps and the same map rotated 180° for maps involved in respective speed combination (“180° Rotation”). Lines join symbols representing individual neurons (FA-1 – blue, SA-1 – red) and bars indicate mean values across all neurons.

Despite the fact that between the mappings at the different speeds, a neuron was subjected to 15 scans involving another pattern of raised elements causing generation of several thousands of action potentials (see Methods), a neuron’s maps obtained at the different scanning speeds were strikingly similar. Averaged across all 16 neurons, the correlation (R^2^) was 0.83, 0.76 and 0.74 for the following speed combinations 15 and 30 mm/s, 30 and 60 mm/s and 15 and 60 mm/s, respectively (“between speeds” correlation in **Fig. 6B**). Yet, the correlations were somewhat weaker than the correlations between the empirical maps for the speed within the speed-pair that showed the lowest correlation (cf. “between speeds” and “within speed” correlation in **Fig. 6B**). To critically address if a neuron’s subfield layout was preserved across speeds, we investigated whether between-speed correlations within neurons were significantly higher than the mean of correlations obtained with 180° rotation of corresponding maps. Strikingly, for each neuron and all speed combinations the between-speed correlation was distinctly higher (**Fig. 6B**, cf. “Between-speeds” and “180° Rotation”). The between-speed correlation and the correlation involving 180° map rotation was significantly different, which verified this speed invariant characteristics of the sensitivity topography (F_1.14_ = 103.0, P <0.0001). Neither speed combination nor neuron type showed a statistically significant effect on the difference (F_2.28_ = 3.1, P = 0.06; F_1,14_ = 0.41, P = 0.53, respectively) and there was no interaction effect between speed combination and neuron type (F_2,28_ = 0.02, P = 0.98).

Next, we asked how well the distinctiveness a neuron’s receptive field properties across speeds is maintained with reference to other neurons’ fields. For each neuron, we cross-correlated the map obtained at each speed with the neuron’s own maps obtained at the other two speeds and with the maps obtained for all other neurons at each speed (3 speeds x 16 neurons - 1 = 47 correlations/speed). We then assessed for each speed how often the highest and the second highest correlation were found among the same neuron’s maps obtained at another speed. Strikingly, the maps of all 16 neurons and at all three speeds were most similar to a map of the same neuron obtained at one of the other two speeds. Even for just one speed, the probability would be practically zero for this to happen by chance (P = (2/47)^16^). Moreover, for 39 out of the 48 maps, the second-highest correlation was also found with a map of the same neuron, again an outcome that by chance would be virtually zero. We did not find an effect of neuron type on the frequency of cases where the second-highest correlation was with a map of another neuron (Χ^2^_1_ = 1.99, P = 0.158).

Taken together, these results show that a neuron’s receptive field sensitivity topography was largely invariant across tested scanning speeds and that the particularities of the receptive field properties relative to other neurons receptive fields essentially are maintained across speeds. This is in line with previous indications that the spatial structuring of FA-1 and SA-1 responses to scanned raised tactile elements is substantially maintained at speeds up to at least 90 mm/sec (Phillips et al., 1992; Pruszynski and Johansson, 2014).

### Conservation of receptive field sensitivity topography across scanning directions

To examine the stability of the subfield layout across scanning directions, we compared maps generated with 0.1 mm kernel width for scans at 30 mm/s in the proximal-distal and distal-proximal directions. Data from 22 neurons (16 FA-1s, 6 SA-1s) were analyzed, eleven of which (8 FA-1s, 3 SA-1s) were recorded in the present experiment and the remaining eleven (8 FA-1s, 3 SA-1s) in a previous series of experiments (Pruszynski and Johansson, 2014). For the neurons of the present experiments, for each direction the map used was an average of the maps obtained by the three scans, whereas for the remaining neurons only one map was available for each direction. The estimated spatial resolution during scans in the two directions available for the neurons of the present experiment did not differ significantly (t_10_ = 0.68, P = 0.51; T-test for dependent samples).

On visual inspection of a neuron’s maps for the two directions, apparently homologous subfields could usually be identified, but their relative positions in the receptive field could differ between the maps (**Fig. 7A**, top panels). That is, compared to one of the maps, the map of the opposite direction appeared to be subject to different degrees of compression, stretching and shear, and could even appear slightly rotated. Such warping would be consistent with the neurons having ridge associated receptors and that direction-dependent shear deformations of the ridge pattern of varying complexity occur when a surface slides over the fingertip skin (Delhaye et al., 2016).

**Figure 7.**
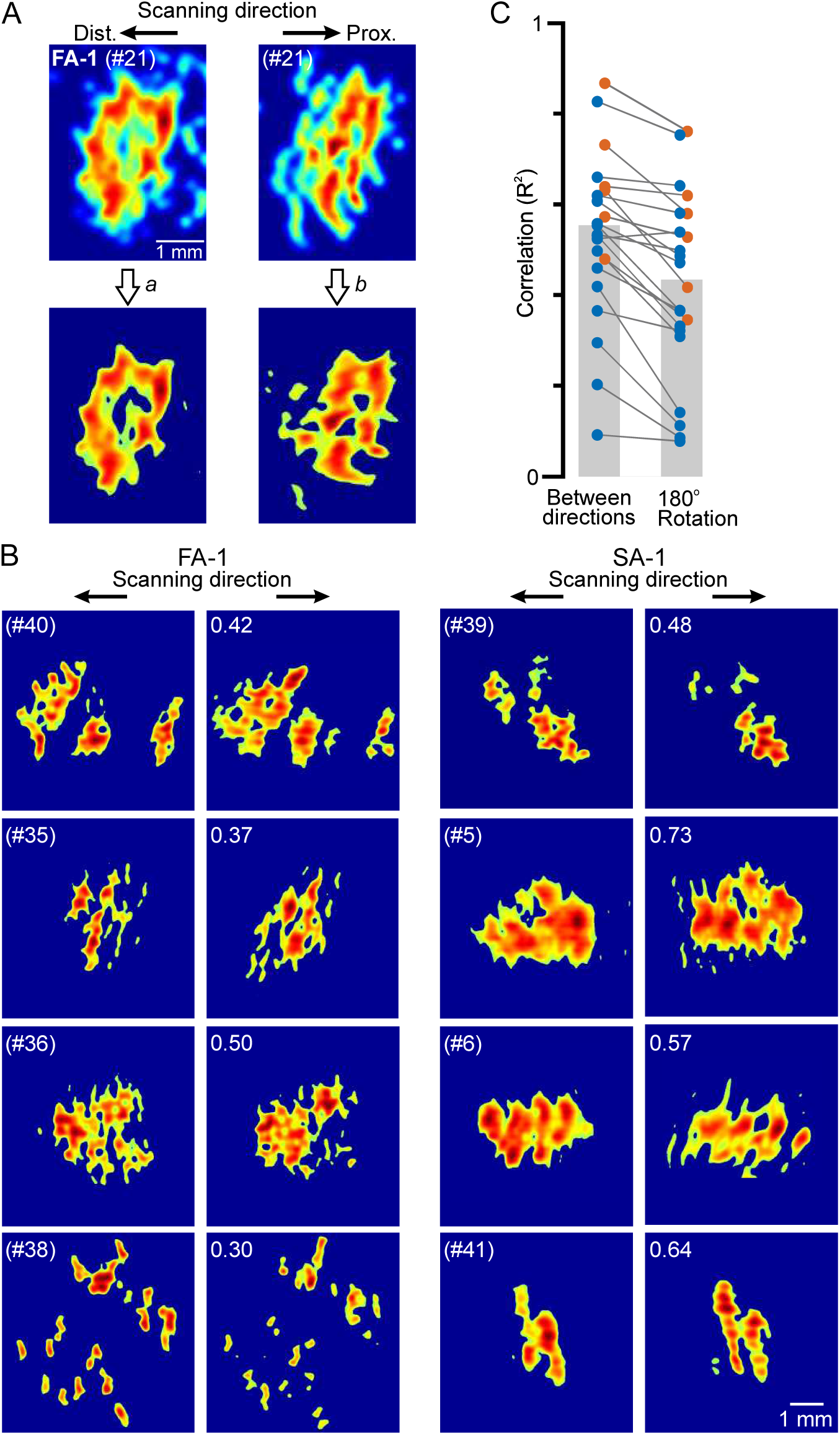
Effect of scanning direction on receptive field sensitivity topography examined at 30 mm/s scanning speed. **A**. Left and right top panels show receptive field sensitivity topography of an exemplar FA-1 neuron obtained in proximal-distal and distal-proximal scanning direction, respectively. Left bottom panel shows the proximal-distal map after thresholding (*a*) and right bottom panel the thresholded distal-proximal map that best matched the thresholded proximal-distal map after transformed (*b*; entire distal-proximal map was stretched in the scanning direction, compressed in its perpendicular direction and counterclockwise rotated). **B**. Comparison of thresholded sensitivity maps of four exemplary neurons of each type obtained during proximal-distal and distal-proximal scanning after the latter had been transformed to best match the former. Numbers in the top left corners of the distal-proximal maps indicate, for each neuron, the correlation between the compared maps. **C**. Pairwise correlations within individual neurons between thresholded proximal-distal and distal-proximal maps after the latter had been transformed (“Between directions”) and between thresholded proximal-distal maps and the same maps rotated 180° (“180° rotation”). Lines join symbols representing individual neurons (16 FA-1 – blue, 6 SA-1 – red) and height of columns indicate mean values across all neurons.

To quantitatively examine the consistency of the subfield layout across the scanning directions in the face of map warping, we performed an analysis where we sought to factor in some aspects of the warping. First, we thresholded the maps to 50% of the maximum value to focus on highly sensitive zones (**Fig. 7A**, *a*). We then transformed the map obtained in the distal-proximal scanning direction to best resemble that obtained in the proximal-distal direction as judged by cross-correlation (**Fig. 7A**, *b*). The parameters of the transformation involved stretching/compressing the entire map both in the scanning direction and in its perpendicular direction and rotation of the map. By changing the values of these parameters with small steps and in different combinations, the coefficients that gave the best correlation were determined and used for the transformation (see Methods). Even though this transformation did not offset shear deformations of the skin surface within the receptive fields, for each neuron type it generally resulted in visually fairly similar maps for the two directions (**Fig. 7B)**. Moreover, the pairwise correlation between a neuron’s maps of the two scanning direction was regularly higher than that between the map of the proximal-distal direction and the same map rotated 180° (F_1,20_ = 38.9, P < 0.0001) (**Fig. 7C**). This indicated that neurons’ subfield structure was reliably preserved over scanning directions.

We finally considered how well a neuron’s receptive field maintains its distinctive character over other neurons’ fields across scanning directions. We cross-correlated each neuron’s processed map with its map for the opposite scanning direction and with the corresponding maps obtained for all other neurons in both directions (2 × 43 correlations). We then evaluated how frequently amongst neurons the highest correlation existed for the same neuron’s maps. Of all 22 neurons we found this happened for 17 and 18 neurons in the proximal-distal and distal-proximal direction, respectively. The chance, at the population level, for this outcome would be virtually zero if the neurons’ distinctiveness regarding receptive field properties would have been lost with the change in scanning direction (P < 0.0001; binominal test).

Taken together, these results suggest that the internal sensitivity topography of a neuron’s receptive field was largely conserved across scanning directions but could be influenced by directional dependent shear deformations of the skin surface. In addition, most neurons retain the distinctiveness of the features of their receptive fields with reference to other neurons’ fields.

## Discussion

Our results indicate that the spatial resolution of the subfield arrangement of FA-1 and SA-1 neurons innervating human fingertips is in the submillimeter range. The estimated spatial resolution as well as the receptive field sensitivity topography appears similar across the tested scanning speeds and the modest speed effect on maximum firing rate indicates that the spatial structuring of neurons’ responses is well maintained even at low tangential speeds. The estimated spatial resolution is also similar across the scanning directions, but the subfields can be displaced relative to one another to some extent depending on the stimulation direction. We interpret this observation as the subfields staying at fixed places on the skin surface and their relative displacement reflecting complex direction-dependent shear deformations of the skin surface as a surface slides over the fingertip, considered to be partially dictated by the fingerprint ridges (Delhaye et al., 2016).

The similarity between the dimensions of the width of the fingerprint ridges and the neurons’ subfields and their estimated spatial resolving power suggests that the ridge-associated receptor organ that accounts for a subfield measures mechanical events at an individual ridge. Such spatial selectivity might be achieved by a combination of the structural compartmentalization of the ridged skin and the ridge-governed contact mechanics of the fingertip. As for the structure, the limiting (adhesive) ridges that anchor the papillary ridges to deeper tissues (Cauna, 1954; Halata, 1975) apparently allow a ridge to be laterally deflected without appreciably affecting its neighbors (Johansson and LaMotte, 1983; LaMotte and Whitehouse, 1986; Lee et al., 2019). Similarly, the transverse ridges protruding into the dermis, mechanically delimiting dermal papillae from each other along a ridge (Cauna, 1954; Halata, 1975), probably explains the spatial selectivity of receptor-organs along a ridge. Regarding contact mechanics, the sliding of the stimulus surface in the current study meant that frictional forces acted on skin ridges, which is almost always the case during object manipulation and tactile exploratory tasks. For smooth parts of the stimulus surface, adhesive frictional forces were likely distributed similarly over microscopic contact zones at the peaks of individual ridges (Soneda and Nakano, 2010; Delhaye et al., 2016) whereas the distinct surface irregularities at the stimulation dots likely caused local phasic distortions of consecutive ridges through interlocking, plowing, and hysteresis friction (Johansson and LaMotte, 1983; LaMotte and Whitehouse, 1986; Tomlinson et al., 2011; Derler and Gerhardt, 2012; van Kuilenburg et al., 2013; Chimata and Schwartz, 2015; Lee et al., 2019). As such, skin deformation caused by sliding tactile stimuli excite primate ridge-associated first-order tactile neurons much more effectively than comparable stimuli perpendicularly indented into the skin (Vallbo and Hagbarth, 1968; Johnson and Lamb, 1981; Phillips et al., 1983; LaMotte and Whitehouse, 1986; Johansson and Westling, 1987). Moreover, the sensitivity topography of FA-1 and SA-1 receptive fields exhibits deeper spatial modulation with sliding stimuli than with punctate perpendicular skin indentations (cf. current results and (Johansson, 1978)). These enhancements in peripheral tactile sensing likely contribute to the increase in perceived intensity and clarity of tactile images and small surface details during sliding movements compared to when we statically contact the same objects.

The current study has several limitations. These include methodological issues that may have resulted in an underestimation of the spatial resolution of neurons’ subfield arrangement. First, mechanical changes in the fingertip with respiration and heartbeats (Johansson and Vallbo, 1979) and varying creep of the skin during the repeated scans (drum revolutions) might have imposed noise in our construction of SEPs and thus falsely increased the spatial jitter of action potentials. Second, the similarity in size of the stimulation dots (top diameter = 0.4 mm) and our estimates of subfield resolution suggest that the probe’s dimension could have acted as a spatial low-pass filter and thus contributed to an underestimation of the resolution.

Technical limitations of our video monitoring system prevented us from recording dynamic interactions between the stimulation dots and papillary ridges with sufficient spatial accuracy and temporal resolution for analysis of direct correlations between individual ridge deformation changes and evoked neural signals. Thus, we could not directly prove that a subfield selectively represented deformation changes of a certain ridge.

Other limitations concern the generalizability of the results. The present and previous functional studies of the subfield arrangement of the FA-1 and SA-1 neurons are based on scanned stimuli limited to ∼ 0.5 mm high embossed elements with trapezoidal cross sections (Phillips et al., 1992; Pruszynski and Johansson, 2014). Thus, little is known about how this arrangement is expressed in responses of FA-1 and SA-1 neurons to scanned geometric stimuli with different heights, curvatures and sharpness etc. Although, effects of such stimulus parameters have been studied in the analogous neurons of monkeys (usually referred to as RA and SA) (LaMotte and Srinivasan, 1987b, a; Lamotte et al., 1994; Blake et al., 1997), the results cannot be translated to humans because their receptive fields do not typically exhibit a corresponding heterogeneous internal sensitivity topography characterized by multiple subfields (Johnson and Lamb, 1981; Phillips and Johnson, 1981; LaMotte and Whitehouse, 1986; Lamotte et al., 1994; Blake et al., 1997; Suresh et al., 2016).

Similarly, the utility of the subfield arrangement of FA-1 and SA-1 neurons in encoding fine tactile texture is unknown. For example, for the FA-1 neurons that are exceptionally sensitive to local skin distortions, the prevailing view based on monkey studies is that they only signal temporal information about vibrations that propagate openly through the skin when exploring fine structures (Phillips and Johnson, 1985; Yoshioka et al., 2001; Weber et al., 2013; Lieber et al., 2017). A possible contribution from a spatial population code comprising spatially modulated patterns of nerve activity based on interactions between texture elements and the neurons’ subfields within the contact area has not been considered. Such a contribution might help to explain a still unsolved problem of fine texture coding, namely how texture perception can be invariant over a wide range of scanning speeds (Weber et al., 2013; Boundy-Singer et al., 2017).

The inability of an individual neuron to signal which of its subfields are primarily stimulated does not preclude the possibility that a population of neurons can signal tactile stimuli at subfield resolution (Pruszynski and Johansson, 2014; Pruszynski et al., 2018; Hay and Pruszynski, 2019). As described previously, the key is that subfields belonging to different neurons are highly intermingled and partially overlap because receptive fields of neurons heavily overlap (Johansson and Vallbo, 1980). Hence, when an object is touched, neurons whose subfields spatially coincide with salient tactile features are primarily excited, while in a slightly different spatial stimulus configuration, another subset of neurons, which can share members with the first subset, is primary excited. Theoretically, for the FA-1 population innervating the fingertips, where all dermal papillae contain Meissner bodies, the resolution of such a spatial coincidence code would approach the distance between adjacent dermal papillae as about half of them are innervated by axonal branches originating from more than one neuron (Matsuoka et al., 1983; Nolano et al., 2003). This scheme would have the added benefit of supporting rapid use of fingertip events in manual dexterity because it could enable “near-instantaneous” coding and processing of spatial tactile stimuli by not requiring integrating signals from individual first-order tactile neurons over time to estimate firing intensity (Johansson and Birznieks, 2004; Johansson and Flanagan, 2009; Pruszynski and Johansson, 2014; Schwarz, 2016; Pruszynski et al., 2018). Indeed, there is substantial evidence that the brain possesses fast feedforward mechanisms for detecting, with millisecond precision, epochs of stimulus-dependent correlations of action potentials in populations of sensory neurons based on relative spike timing (Petersen et al., 2002; VanRullen and Thorpe, 2002; Gollisch and Meister, 2008; Tiesinga et al., 2008; Panzeri and Diamond, 2010; Gutig et al., 2013; Bale et al., 2015; Zuo et al., 2015; Pruszynski and Zylberberg, 2019).

Sampling of spatial tactile patterns with first-order neurons receiving converging inputs from multiple subfields cannot allow for complete reconstruction of all arbitrary patterns at subfield resolution. However, given the sparsity in biologically relevant signaling patterns, functional spatial acuity corresponding to the subfield resolution could be achieved by the brain by learning mechanisms and localized random sampling as has been described for sensory systems generally (Olshausen and Field, 2004; Barranca et al., 2014; Yamins et al., 2014; Rongala et al., 2018; Zhao et al., 2018). Hence, the sub-field organization may support functional spatial acuity corresponding to the density of the receptor organs in the skin but with far fewer axons in the peripheral nerves than if the receptors were innervated separately. Dealing with fewer input channels might also contribute to increased central neural process speed.

## Acknowledgments

This work was supported by the Swedish Research Council (Projects 22209 and 01635). J.A.P was funded by a Long-Term Fellowship from the Human Frontier Research Council. We wish to thank Carola Hjältén, Göran Westling, Anders Bäckström and Per Utsi for technical support.

## Notes

### Competing Interest Statement

The authors have declared no competing interest.

